# Chronic hyperglycemia drives alterations in macrophage effector function in pulmonary tuberculosis

**DOI:** 10.1101/2022.01.18.476721

**Authors:** Sudhasini Panda, Diravya M Seelan, Shah Faisal, Kalpana Luthra, Jayanth Kumar Palanichamy, Anant Mohan, Naval K Vikram, Neeraj Kumar Gupta, Lakshmy Ramakrishnan, Archana Singh

## Abstract

**Background:** The rising prevalence of Diabetes mellitus (DM) in high TB endemic countries has the potential to adversely affect sustainability of TB control since DM can lead to alterations in both innate and adaptive immune response constituting as a risk factor for development of active tuberculosis (TB). The impact of hyperglycemia on TB specific innate immune response in terms of macrophage functions remains poorly addressed.

**Material and methods:** Macrophage effector functions in diabetic and non-diabetic individuals with and without PTB infection as well as non-diabetic-uninfected controls (fifty individuals in each group) were assessed. Phagocytic capacity against BCG and surface expression of PRRs (CD11b, CD14, CD206, MARCO and TLR2) were measured via flow cytometry. Effector molecules (ROS and NO) were assessed via DCFDA and Griess reaction respectively.

**Results:** A systematic dysregulation in phagocytic capacity with concurrent alterations in expression pattern of key PRRs (CD11b, MARCO and CD206) and effector molecules (ROS and NO) was observed in diabetic individuals with PTB. These altered macrophage functions were positively correlated with increase in disease severity in diabetic individuals.

**Conclusion:** Our results highlight several key patterns of immune dysregulation against *M.Tb* under hyperglycemic conditions. A significant reduction in macrophage effector functions in infected diabetic individuals which further correlated with increase in disease severity reveals a negative impact of hyperglycemia with aetiology and pathological progression of TB.

## Introduction

With WHO 2019 report labelling diabetes and TB as co-epidemic and, increased incidence of TB in diabetes have led to renewed interest in understanding pathophysiology of this co-epidemic (1). There is an urgent need to implement strategies for TB prevention among the millions of DM patients exposed to *Mycobacterium tuberculosis* (*M.Tb*) worldwide, but knowledge is limited on how and when DM alters the natural history of this infection. The increased incidence of TB in people with DM appears to be multifactorial. Recent evidence has shown that DM patients tend to have a higher incidence, higher disease severity, including development of drug-resistance TB, compared with those without DM. Chronic DM is associated with delayed innate immunity to *M.Tb.* due to late delivery of *M.Tb*-bearing antigen-presenting cells to the lung draining lymph nodes (2). Efficient phagocytosis and priming of the adaptive immune response are necessary to activate the cell-mediated immune responses that restrict initial *M.Tb.* growth and these delays likely contribute to the higher risk of DM patients for development of *M.Tb.* infection and persistence (3). Recent evidences suggest that innate as well as adaptive immune responses might be affected (4). Few mice model studies have suggested defect in immune response in terms of immune cell recruitment at site of infection and increased bacterial load in diabetic mice as compared to euglycemic mice (2,5). However, limited in vivo data is available in context of TB-DM. In this regard, understanding of alterations or disturbances in innate immune response in diabetic patients having TB infection is needed.

As the pathogen enters the host, it is recognized by innate cells like macrophages via pathogen recognition receptors followed by the phagocytosis of the pathogen and ultimately killing via formation of reactive oxygen and nitrogen species (6). Macrophages are key to the aetiology of TB due to their dual role as a primary host cell reservoir for M. Tb, as well as being effector cells that control and eliminate *M.Tb.* (7,8). The examination of alveolar macrophages in TB-DM patients has revealed the presence of hypodense alveolar macrophages, which are less activated and are correlated with the severity of disease, implying that they might contribute to the increased susceptibility to *M.Tb.* infection (9). Little is known about the state of macrophage activation during the low-grade chronic inflammation linked to diabetes mellitus. Several pathogen recognition receptors (PRRs) like TLR 2, Mannose receptor (CD206), Complement receptor 3, Macrophage receptor with collagenous structure (MARCO) and CD14 are involved in recognition of the *Mycobacterium tuberculosis* bacillus (10). After recognition, the bacteria are phagocytosed by the immune cells mainly by macrophages followed by phagocytic killing via nitric oxide (NO) and reactive oxygen species (ROS) (11). Chronic inflammatory conditions like diabetes mellitus also lead to production of ROS (12). Overstimulated ROS production under this condition may modulates the inflammatory network in TB infection, leading to dysfunctional inflammatory responses and tissue remodelling having adverse effect on host.

Taking this available literature under consideration, we hypothesized an increased susceptibility of diabetic patients to TB could be due to the defects in macrophage effector functions like bacterial recognition, phagocytic activity, killing via reactive oxygen and nitrogen species and cellular activation due to hyperglycemia which could result in impaired or dysregulated immune response. Impaired immune response and killing of intracellular bacteria will then potentially increase bacterial load, chronic inflammation, and central necrosis that would facilitate bacterial dissemination. Limited human studies are available in this area and none in diabetic patients having TB (PTB + DM) cohort.

The present study was designed to assess the alterations in macrophage effector function in terms of bacterial recognition followed by phagocytosis and bacterial killing via ROS/RNS in diabetic patients having TB. Surface expression of different PRRs like TLR 2, Mannose receptor (CD206), Complement receptor 3, Macrophage receptor with collagenous structure (MARCO) and CD14 receptors in conjunction with functional properties such as phagocytosis, ROS and NO production, were studied in DM patients having TB. Understanding of defects in innate immune responses in diabetic condition could help in early identification of the active disease among diabetic individuals and future development of new treatment targets to limit development of TB among them.

## Material and Methods

### Study subjects

The cross-sectional study was conducted in Department of Biochemistry and Department of Medicine, All India Institute of Medical Sciences, New Delhi. Study participants were recruited under four groups namely pulmonary TB group (PTB), type 2 diabetes mellitus (DM) group, PTB + DM group and healthy control group.

65 newly diagnosed treatment naïve pulmonary TB patients (PTB) were enrolled under PTB group. A PTB case was defined as a patient with clinically diagnosed case of TB affecting the lungs, having symptoms of fever or cough and sputum smear that showed acid-fast bacilli or culture positive for *M.Tb.* or Gene Xpert. Any other conditions like extra-pulmonary and drug resistant tuberculosis, HIV, diabetes, hypertension, significant organ dysfunction of heart, liver and kidney, abnormal hematologic function, inflammation like autoimmune disease, atopic dermatitis, pregnant or lactating women, or any other uncontrolled concurrent illness were excluded.

51 uncontrolled type 2 diabetic patients with HbA1c levels >7.5 (DM) without any other disorders as mentioned above including tuberculosis were recruited under diabetes (DM) group. Individuals taking metformin, corticosteroids, aspirin or TNF blockers were also excluded to avoid the confounding factors which may alter the immune function.

50 newly diagnosed treatment naïve pulmonary TB patients who had uncontrolled type 2 diabetes mellitus with HbA1c > 7.5% (PTB + DM) were also recruited for the study with above mentioned inclusion and exclusion criteria.

50 individuals with no known history of TB and diabetes or any other disorders were recruited as healthy controls. All study participants were age (18-50 years) and sex matched. Height and weight were recorded to calculate body mass index. The study was approved by institutional ethics committee (IECPG-374/28.09.2017). Study participants were recruited from north Indian population after obtaining informed consent.

### Culture of *M. bovis* BCG and labelling with FITC

*M. bovis* BCG was used and grown to the log phase in 7H9 middlebrook medium supplemented with oleic albumin dextrose catalase (OADC). The bacteria were then harvested, washed, and frozen at −80°C in PBS plus 10% of glycerol. Bacterial load was determined by plating serial 10-fold dilutions on 7H10 middlebrook agar (supplemented with OADC (M0678)) and counting colonies after incubation for at least 3 weeks. ZN staining was performed to rule out contamination by other bacteria (Supplementary figure 1).

### Generation of monocyte derived macrophages (MDMs)

PBMCs were isolated from whole blood using Ficoll gradient separation. For monocyte isolation by plastic adherence method, 1-2 million PBMCs were plated into 12 well plate (Nunc, Thermo Fisher) at 37°C and 5% CO_2_ and allowed to adhere for two hours in 1 mL of RPMI 1640 (Hyclone) supplemented with 10% fetal bovine serum (Himedia) and 1 % penicillin/streptomycin. After two hours, non-adherent cells were removed and adhered monocytes were then cultured into macrophages using 35 ng/ml GM-CSF for 9 days along with addition of autologous serum at every third day. After 9 days, cells were differentiated into macrophages with larger size and protruding appendages (Supplementary figure 2). The purity was checked by using CD11b and CD14 by flow cytometry using flurochrome tagged antibodies [PE-Cy7-CD11b (557743) and PerCP-Cy5.5-CD14 (562692)]. Cells with CD11b^high^ CD14^low^ were taken as differentiated macrophages (Supplementary figure 3). The cell viability was checked by trypan blue and more than 90% cells were found to be viable. Differentiated macrophages were then taken forward for further experiments.

### Phagocytic activity of macrophages using FITC labelled BCG

Phagocytosis activity of differentiated macrophages were studied using fluorescent detection method. Briefly, BCG were tagged with FITC by incubating 0.2 μl of 5mg/ml FITC for one hour at room temperature with end-to-end rotation followed by washing with PBS to remove unbound FITC. Macrophages were then incubated with FITC labelled BCG at MOI of 10 for one hour at 37°C. After incubation, 1 ml of ice cold 1X PBS was added next to stop phagocytosis. Cells were scraped out and washed with 1X PBS (3 times) to remove free bacteria. Cells were then stained with CD11b and CD14 for macrophage purity. To distinguish cells which have phagocytosed bacteria from those simply binding the bacteria at the surface, 100 μl of trypan blue was added to the cells and incubated for one minute to quench surface FITC fluorescence. Cells with phagocytosed bacteria was then analyzed using flow cytometry and median fluorescence intensity (MFI) values and percentage positivity were recorded.

### Surface expression of different pathogen recognition receptors on macrophages using flow cytometry

Surface expression of different pathogen recognition receptors namely TLR2, Mannose receptor (CD206), Complement receptor 3, MARCO and CD 14 were studied on macrophages by flow cytometry using fluorochrome tagged antibodies [PE mouse anti-human TLR 2 (565349), APC mouse anti-human CD206 (550889), Mouse anti-human MARCO antibody (HM2208) + FITC conjugated goat anti-mouse secondary antibody (Ab97259), PE-Cy7-CD11b (557743), PerCP-Cy5.5-CD14 (562692)]. Briefly 1×10^5^ macrophages were stained with titrated volume of antibody cocktail for 30 minutes at 37°C. Cells were then washed with FACS buffer (PBS + 0.5% BSA). After staining, cells were acquired on BD LSR fortessa X-20 and median fluorescence intensity (MFI) values and percentage positivity were recorded. Analysis was performed on FlowJo V10. FMO (fluorescence minus one) was used for gating the signals.

### Levels of Reactive oxygen species (ROS) by flow cytometry and serum nitric oxide (NO) levels by griess reaction

Reactive oxygen species in macrophages were estimated using Dichloro-dihydro-fluorescein diacetate (DCF-DA) assay (ab113851). Briefly, 1-2 × 10^4^ macrophages were taken in flow tube. Then 10^5^ bacteria were added to the cells 1 hour prior to the treatment. 500ul of DCFDA solution was added at a concentration of 20 uM followed by 30 min incubation at 37°C. Signal was read using flow cytometry.

NO levels were measured in serum using colorimetric assay based on griess reaction. Griess reagent includes naphthylethylenediamine dihydrochloride suspended in water and sulphanilamide in phosphoric acid. This reagent reacts with nitrite in samples to form a purple azo product, absorbance of which is measured at 540 nm. Serum and Griess reagent was added in 1:1ratio. The mixture was incubated for 10 minutes followed by reading absorbance at 540 nm. The concentration of nitrite in the sample was determined from a sodium nitrite (NaNO_2_) standard curve

### Statistical Analysis

All statistical analyses were performed on GraphPad Prism 6 (GraphPad Software Inc., San Diego, CA, USA) or R statistical software version 4.1.1 (R Project for Statistical Computing) within RStudio statistical software version 1.4.1717 (R Studio). Categorical variables were presented as counts and percentages. Continuous variables were reported as mean (SD) or median (interquartile range [IQR]) after assessment of normality. Non-parametric statistical analyses were performed throughout the study after checking for normality. Correlation between variables were assessed using Spearman correlation. Mann whitney U and Kruskal wallis test were used for comparison between two groups and three groups respectively. Linear regression analyses were used to determine factors (PRRs) associated with phagocytosis and were reported as beta coefficients and 95% CIs. A 2-tailed p < 0.05 was considered statistically significant for all conducted analyses.

## Results

### Clinical and laboratory data of study participants

The present study included a total of 221 study participants who were recruited under four groups namely pulmonary tuberculosis patients (PTB), uncontrolled Type 2 diabetes mellitus patients (DM), pulmonary tuberculosis patients having uncontrolled type 2 diabetes (PTB + DM) and healthy controls based upon the inclusion and exclusion criteria mentioned in the methodology section. The demographic and clinical characteristics are shown in Table 1.

**Table 1.**
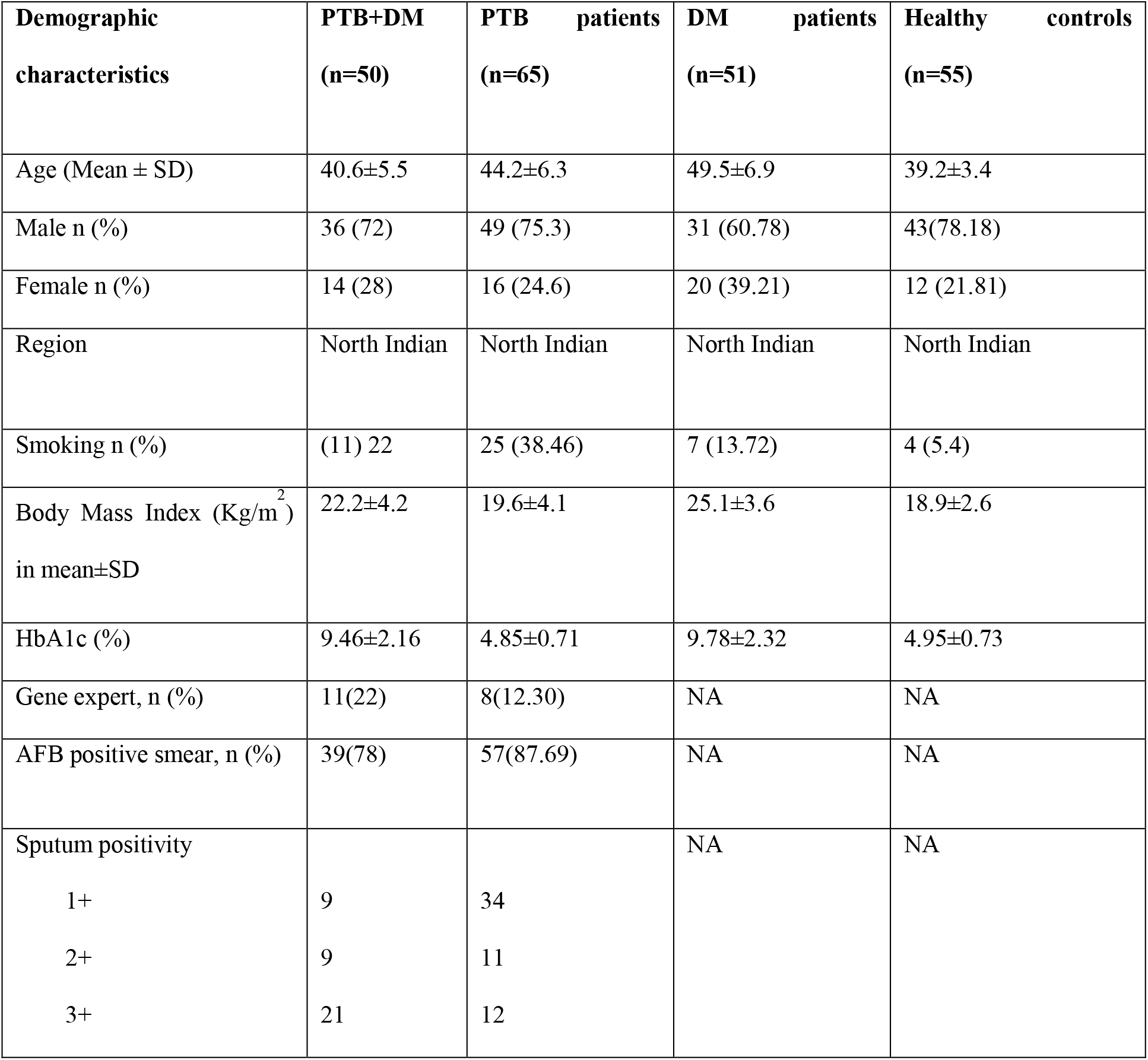
Demographic and clinical characteristics of study participants

All recruited pulmonary TB patients with or without type 2 DM underwent sputum AFB test or gene xpert test as confirmatory test for tuberculosis. Among 65 PTB patients, 87.69% were sputum positive, out of which 59.6% were 1+ grade, 19.29% were 2+ grade and 21.05% were 3+ grade. However, in PTB + DM group, 78% were sputum positive, out of which the frequency of 3+ grade was higher (53.84%) as compared to 2+ (23.07%) and 3+ (23.07%). Since the phenotype of PTB + DMs might differ between study participants who were diabetic prior to TB incident and those with an initial diagnosis of diabetes at the time of PTB incident, we restricted the recruitment criteria to participants with pre-existing uncontrolled diabetes with initial diagnosis of PTB.

### Chronic hyperglycemia variably affects phagocytosis capacity of monocyte derived macrophages (MDMs)

To determine if chronic hyperglycemic state alters macrophage functions, differentiated macrophages were evaluated for the degree of phagocytosis of FITC labelled BCG. Phagocytic capacity was found to be higher in all patient groups as compared to healthy controls as shown in figure 1a and 1b (p<0.0001). In between patient groups, phagocytosis capacity was found to be significantly decreased in PTB + DM and DM patients compared to PTB patients (p< 0.001) suggesting defect in phagocytic capacity of macrophages under chronic hyperglycemic milieu. PTB + DM and DM group had comparable phagocytosis capacity.

**Figure 1.**
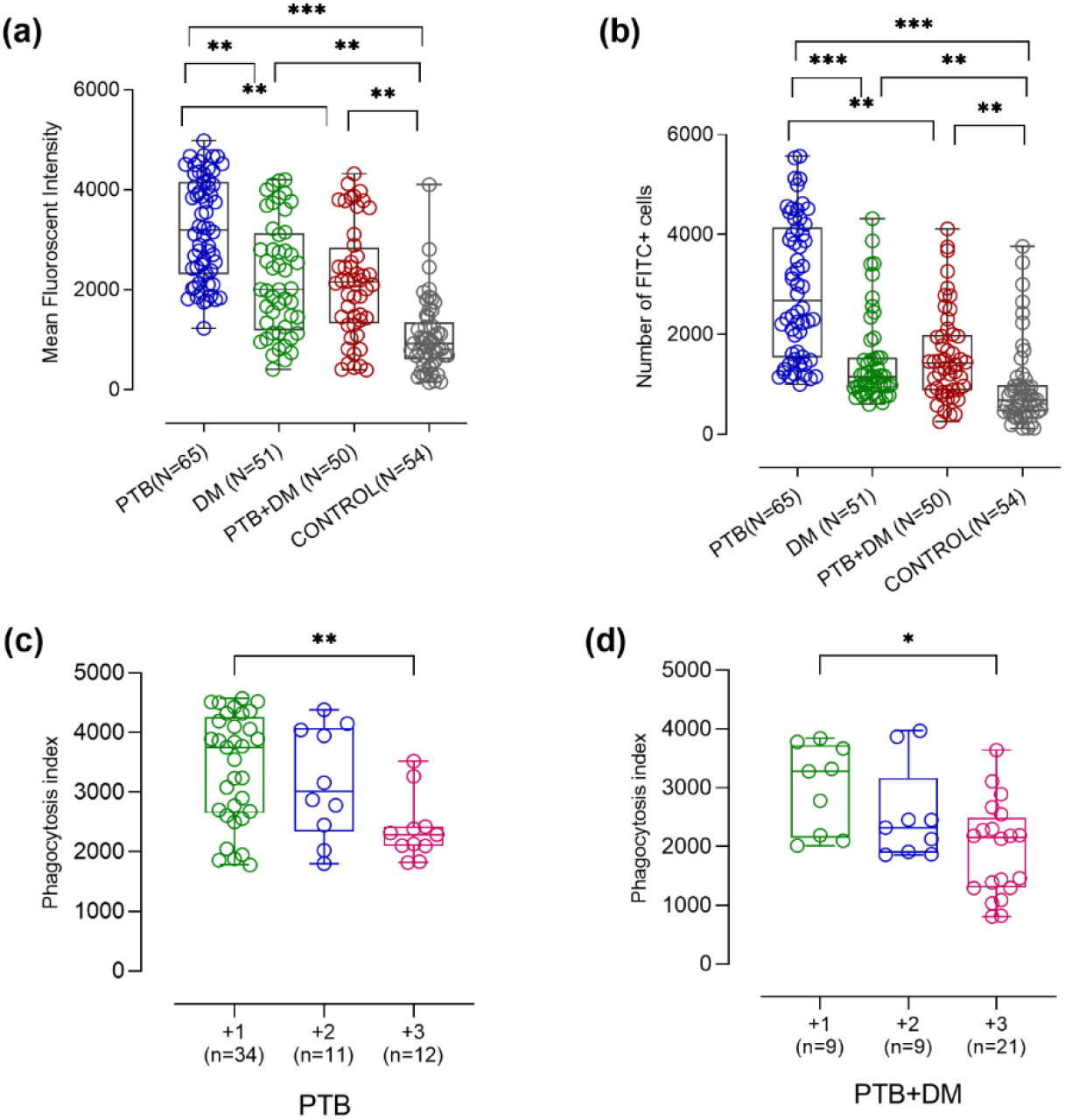
Phagocytosis capacity of monocyte derived macrophages of study participants. 2a represents phagocytosis capacity of cultured macrophages after BCG infection in all study groups namely PTB, DM, PTB+DM and controls. 2b and 2b represents phagocytosis of cultured macrophages after BCG infection in different sputum grade PTB and PTB+DM patients. Macrophages were incubated with FITC labelled BCG and phagocytosed bacteria was visualized using flow cytometry. Data is represented as fluorescent intensity and each point represents individual sample value. Box plot represents median with interquartile range. Kruskal-Wallis testing with post-hoc Dunn’s multiple comparison testing was performed. p values < 0.05 were considered to be statistically significant. One asterisk (*) indicates a p-value < 0.05; two asterisks (**) indicate a p-value < 0.01, three asterisks (***) indicate a p-value < 0.001 and four asterisks (****) indicate a p-value < 0.0001. PTB = Naïve active pulmonary TB; DM= Uncontrolled diabetic patients, PTB+DM= Uncontrolled diabetic patients with pulmonary TB, control= Healthy controls with no history of TB and DM. 1+ = 1+ sputum positive PTB/PTB+DM; 2+ = 2+ sputum positive PTB/PTB+DM; 3+ = 3+ sputum positive PTB/PTB+DM.

PTB and PTB + DM patients were subdivided into different sputum grades to check for alterations in phagocytosis capacity of macrophages with respect to disease severity. We observed significantly decreased phagocytosis capacity with increased sputum positivity or increased disease severity in both PTB and PTB + DM patients (p<0.007 and 0.02 respectively) as shown in figure 1c and 1d.

### Chronic hyperglycemia leads to altered expression of different PRRs on MDMs

Surface expression of different PRRs namely CD11b, CD14, MARCO, TLR2 and CD206) was significantly altered in disease groups compared to healthy controls (figure 2a-2e). Complement receptor 3 or CD11b was found to be significantly decreased in PTB + DM patients as compared to PTB patients (p<0.01). The levels were comparable in DM and PTB + DM patients. CD14 levels were comparable between PTB + DM and DM patients, however, slightly higher than PTB patients (p<0.05). MARCO levels were significantly decreased in PTB + DM patients compared to PTB (p<0.05). The levels of TLR 2 was found to be higher in all patient group as compared to controls (p<0.05). However, levels were comparable between PTB, DM and PTB + DM. CD206 levels were found to be significantly higher in DM milieu in both PTB + DM and DM patients as compared to PTB (p<0.01 and 0.001), suggesting defect in bacterial uptake under chronic hyperglycemic milieu due to alterations in important pathogen recognition receptors.

**Figure 2.**
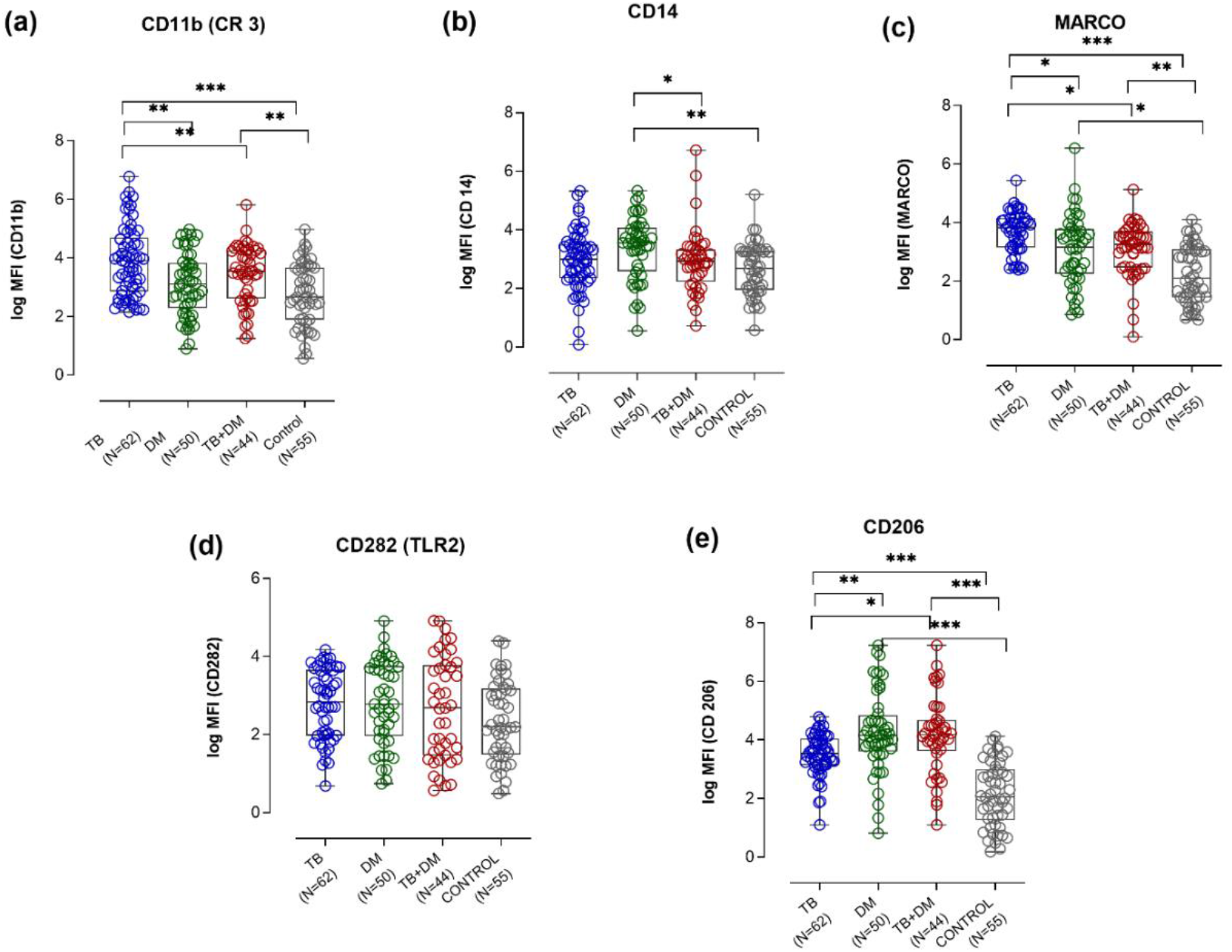
Surface expression of different pathogen recognition receptors on macrophages of study groups namely PTB, DM, PTB+DM and controls. Surface expression was checked by flow cytometry using specific fluorochrome tagged antibodies (a) shows MFI for CD11b, (b) shows MFI for CD14, (c) shows MFI of MARCO, (d) shows MFI of TLR 2, (e) shows MFI of CD206. Data is represented as fluorescent intensity and each point represents individual sample value. Box plot represents median with interquartile range. Kruskal-Wallis testing with post-hoc Dunn’s multiple comparison testing was performed to determine whether expression was statistically different among the different study group. p values < 0.05 were considered to be statistically significant. One asterisk (*) indicates a p-value < 0.05; two asterisks (**) indicate a p-value < 0.01, three asterisks (***) indicate a p-value < 0.001 and four asterisks (****) indicate a p-value < 0.0001. PTB = Naïve active pulmonary TB; DM= Uncontrolled diabetic patients, PTB+DM= Uncontrolled diabetic patients with pulmonary TB, control= Healthy controls with no history of TB and DM

Since we found significant difference in levels of different PRRs in our patient group, we tried to evaluate whether there is any correlation with disease severity. Therefore, we subdivided PTB and PTB + DM patients based upon their sputum positivity into 1+, 2+ and 3+ sputum positive patients and assessed the levels of CD11b, CD14, MARCO, CD206 and TLR2 as shown in figure 3a-3e and 3f-3j respectively. We observed significantly decreased levels of CD11b levels in 3+ sputum positive patients as compared to 2+ and 1+ sputum positive patients of both PTB and PTB + DM patients (p<0.01 and 0.06 respectively). In case of CD14, no difference was observed in different sputum positive PTB patients (p<0.11). However, the levels were significantly decreased in 3+ sputum positive PTB + DM patients as compared to 1+ patients (p<0.02). A similar trend of lower expression levels with increased disease severity was observed for MARCO in both PTB and PTB + DM patients with lowest levels seen in 3+ sputum positive patients (p<0.0001 and 0.001 respectively). Furthermore, levels of TLR2 were significantly decreased in 3+ patients as compared to 1+ and 2+ in PTB + DM patients (p<0.0006). However, no difference was found in PTB patients (p<0.25). Similarly, no significant difference was observed in CD206 levels of different sputum positive patients of PTB group. However, significantly higher levels of CD206 were observed in 3+ sputum positive PTB + DM patients as compared to 1+ and 2+ sputum positive patients (p<0.04, altogether suggesting altered expression of these receptors affecting disease severity.

**Figure 3.**
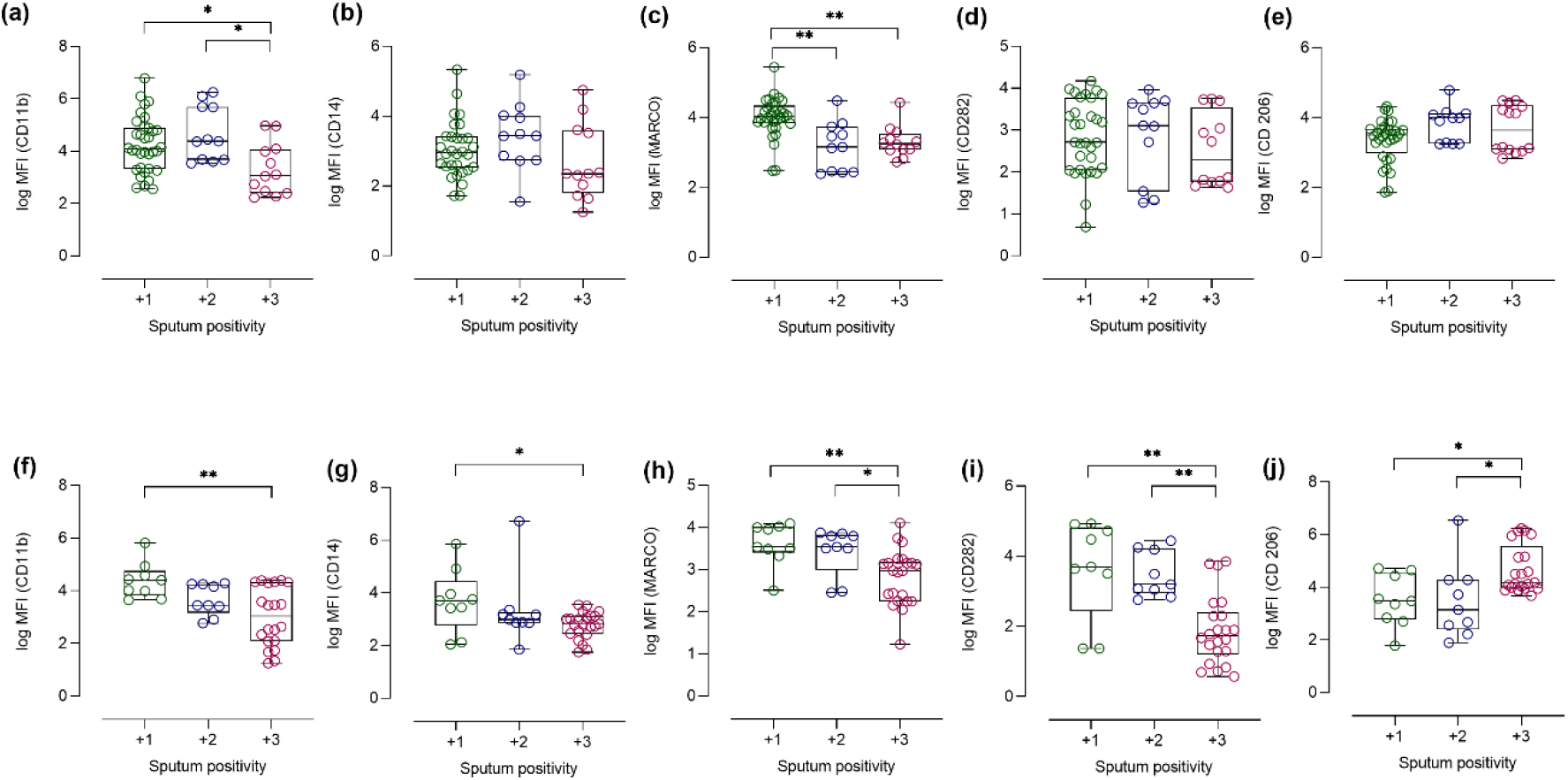
Surface expression of different pathogen recognition receptors on macrophages of different sputum grade PTB and PTB+DM patients. 5a-e shows MFI for CD11b, CD14, MARCO, TLR 2 and CD206 in PTB patients. 5f-j shows MFI for CD11b, CD14, MARCO, TLR 2 and CD206 in PTB + DM patients. Data is represented as fluorescent intensity and each point represents individual sample value. Box plot represents median with interquartile range. Kruskal-Wallis testing with post-hoc Dunn’s multiple comparison testing was performed to determine whether expression was statistically different among the different study group. p values < 0.05 were considered to be statistically significant. One asterisk (*) indicates a p-value < 0.05; two asterisks (**) indicate a p-value < 0.01, three asterisks (***) indicate a p-value < 0.001 and four asterisks (****) indicate a p-value < 0.0001. 1+ = 1+ sputum positive PTB/PTB+DM; 2+ = 2+ sputum positive PTB/PTB+DM; 3+ = 3+ sputum positive PTB/PTB+DM.

### Correlation of CD14 with its coreceptors, MARCO and TLR 2 in monocyte derived macrophages (MDMs) of PTB, DM, PTB + DM and controls

Since MARCO and CD14 act as heterodimer to exert their function, we next assessed the correlation of surface expression of MARCO and CD14 present on macrophages of all the study groups. We found positive correlation in levels of MARCO and CD14 in PTB and healthy control group (r = 0.71 and 0.76). However, no correlation was found in PTB + DM and DM group as shown in figure 4a-4d suggesting defect in bacterial uptake via CD 14-MARCO in diabetic milieu.

**Figure 4.**
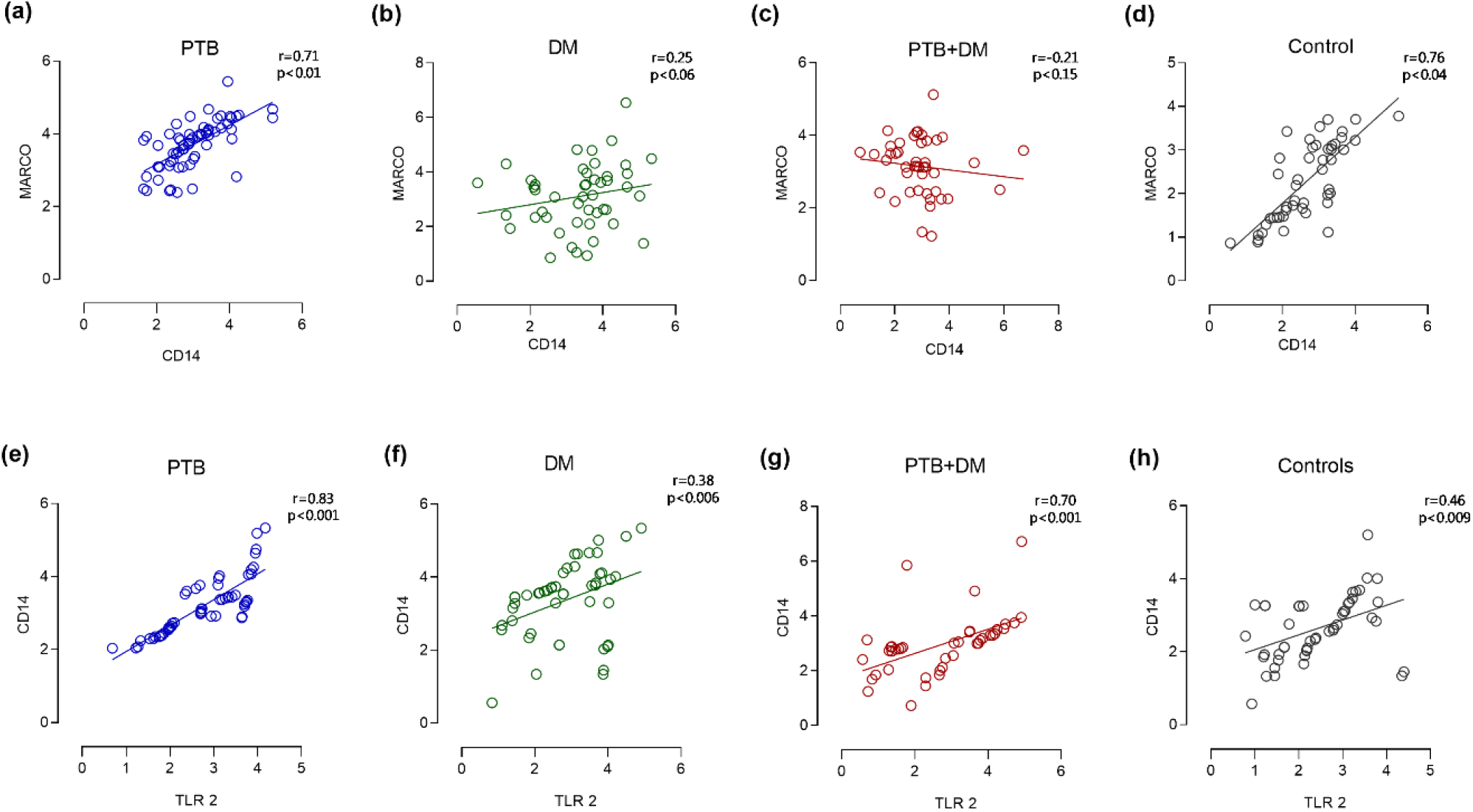
Correlation of CD 14 with MARCO and TLR 2 in all study groups. 4a-d shows correlation between surface expression of MARCO and CD14 on macrophages of all the study groups namely PTB, DM, PTB+DM and controls respectively. 4e-h shows correlation between surface expression of TLR 2 and CD14 on macrophages of all the study groups namely PTB, DM, PTB+DM and controls respectively. Non parametric spearman correlation was used correlation analysis. Spearman’s correlation coefficient is displayed on the right. PTB = Naïve active pulmonary TB; DM= Uncontrolled diabetic patients, PTB+DM= Uncontrolled diabetic patients with pulmonary TB, control= Healthy controls with no history of TB and DM.

Along with MARCO, CD14 is also required for TLR 2 activation and initiation of downstream signalling pathway. Therefore, we correlated the surface expression of TLR 2 and CD14 present on macrophages of all the study groups. We found positive correlation in levels of TLR 2 and CD14 in PTB and PTB + DM group (r = 0.72 and 0.69 respectively). However, no correlation was found in DM and healthy control group as shown in figure 4e-4h.

### Phagocytic capacity is associated with altered PRRs expression on MDMs under chronic hyperglycemia

Phagocytosis capacity of macrophages can be affected by the surface expression of different pathogen recognition receptors, therefore, the association of these PRRs (independent variable) with phagocytosis (dependant variable) was evaluated using multiple linear regression in all the study groups. Phagocytosis capacity was associated with the PRRs namely CD11b, CD14 and CD206 in PTB + DM group (Table 2). Upon correlation analysis, in PTB + DM group, CD11b expression was found to be positively correlated with MARCO and TLR 2 while CD206 was negatively correlated with other receptors as shown in correlation matrix (figure 5a). A similar pattern was observed in DM group with weak correlation as shown in correlation matrix (figure 5b). In PTB group, CD11b, MARCO and TLR 2 were positively correlated among each other as shown in correlation matrix (figure 5c). A weak correlation was found among these PRRs in healthy control group (figure 5d). Similarly, phagocytosis index was also found to be positively correlated with CD11b, MARCO and TLR 2 and negatively correlated with CD206 in patient group (figure 5a-5d).

**Table 2.** Variables (pathogen recognition receptors) associated with phagocytosis in study groups

| **Parameter estimates** | **Variable** | **Estimate** | **P value** | **P value summary** | **95% confidence interval** |
| --- | --- | --- | --- | --- | --- |
| **PTB + DM** |  |  |  |  |  |
| β1 | CD11b | 303.5 | 0.0172 | * | 56.45 to 550.6 |
| β2 | CD14 | 168.5 | 0.0409 | * | 7.282 to 329.7 |
| $\beta_3$ | MARCO | 216.9 | 0.1629 | ns | -91.12 to 524.9 |
| $\beta_4$ | TLR 2 | 118.1 | 0.1902 | ns | -60.77 to 296.9 |
| $\beta_5$ | CD206 | -336.7 | 0.0022 | ** | -545.1 to -128.4 |
| **PTB** |  |  |  |  |  |
| $\beta_1$ | CD11b | 285.8 | 0.0083 | ** | 76.45 to 495.2 |
| $\beta_2$ | CD14 | 89.09 | 0.4543 | ns | -147.6 to 325.8 |
| $\beta_3$ | MARCO | 411.4 | 0.0095 | ** | 104.1 to 718.7 |
| $\beta_4$ | TLR 2 | -6.721 | 0.9604 | ns | -276.8 to 263.3 |
| $\beta_5$ | CD206 | 113.5 | 0.4387 | ns | -177.8 to 404.8 |
| **DM** |  |  |  |  |  |
| $\beta_1$ | CD11b | 294.4 | 0.0183 | * | 52.17 to 536.5 |
| $\beta_2$ | CD14 | -124.2 | 0.2327 | ns | -331.0 to 82.58 |
| $\beta_3$ | MARCO | 21.25 | 0.8217 | ns | -167.6 to 210.1 |
| $\beta_4$ | TLR 2 | 125.6 | 0.2274 | ns | -81.13 to 332.4 |
| $\beta_5$ | CD206 | -464.2 | 0.001 | *** | -673.9 to -254.5 |
| **Controls** |  |  |  |  |  |
| $\beta_1$ | CD11b | 183.1 | 0.0981 | ns | -35.21 to 401.4 |
| $\beta_2$ | CD14 | 46.90 | 0.7062 | ns | -202.0 to 295.8 |
| $\beta_3$ | MARCO | 143.4 | 0.1932 | ns | -75.22 to 362.1 |
| $\beta_4$ | TLR 2 | 1.942 | 0.9872 | ns | -239.9 to 243.7 |
| $\beta_5$ | CD206 | 111.9 | 0.2507 | ns | -81.72 to 305.5 |

Multiple linear regression analysis for association of phagocytosis (dependant variable) with different PRRs (independent variables) namely CD11b, CD14, MARCO, TLR 2 and CD206 in all study groups. Data is represented as 95% Confidence Interval. P<0.05 was considered significant

**Figure 5.**
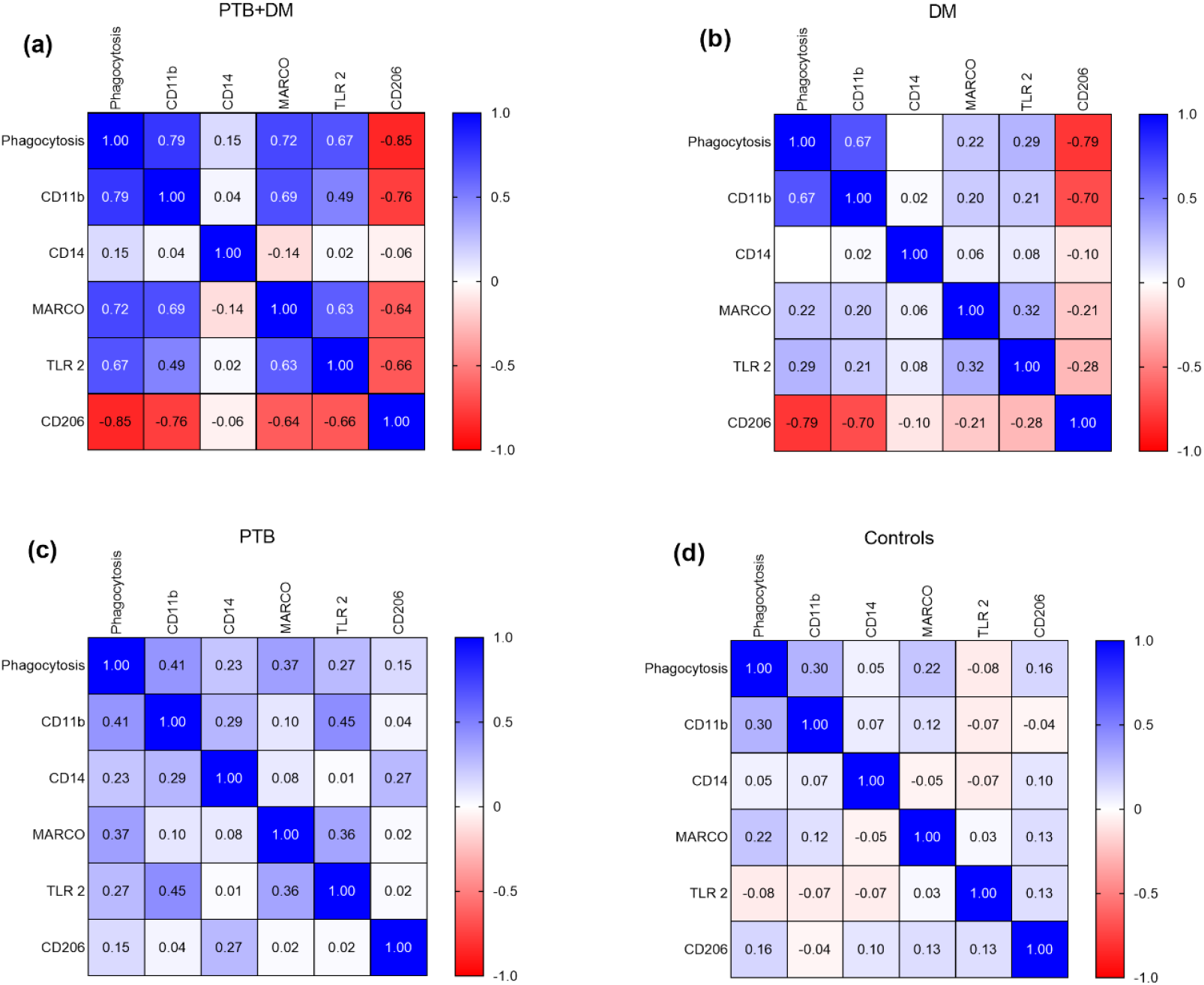
Correlation matrix surface expression of PRRs and phagocytosis index of macrophages in study groups. 5a-d shows values of PTB+DM, DM, PTB and control group respectively. Spearman correlation was used to correlate the variables. PTB+DM= Uncontrolled diabetic patients with pulmonary TB. Values in each block shows Spearman’s correlation coefficient.

### Chronic hyperglycemia drives aberrant production of reactive oxygen species and nitric oxide

We have observed that ROS levels were significantly higher in PTB patients compared to healthy controls (p<0.001) due to ongoing infection in these individuals. However, ROS levels were even higher in DM and PTB + DM patients as compared to PTB patients (p<0.01 and 0.001 respectively) and healthy controls (p<0.001) as shown in figure 6a suggesting overstimulation of ROS production under chronic hyperglycemic milieu. While comparing ROS levels in different sputum grade patients, no significant difference was observed in ROS levels of different sputum positive PTB patients (p<0.08). However, in PTB + DM patient group, higher levels of ROS were observed in 3+ sputum positive patients as compared to 2+ and 1+ (p<0.005) as shown in figure 6b-6c.

**Figure 6.**
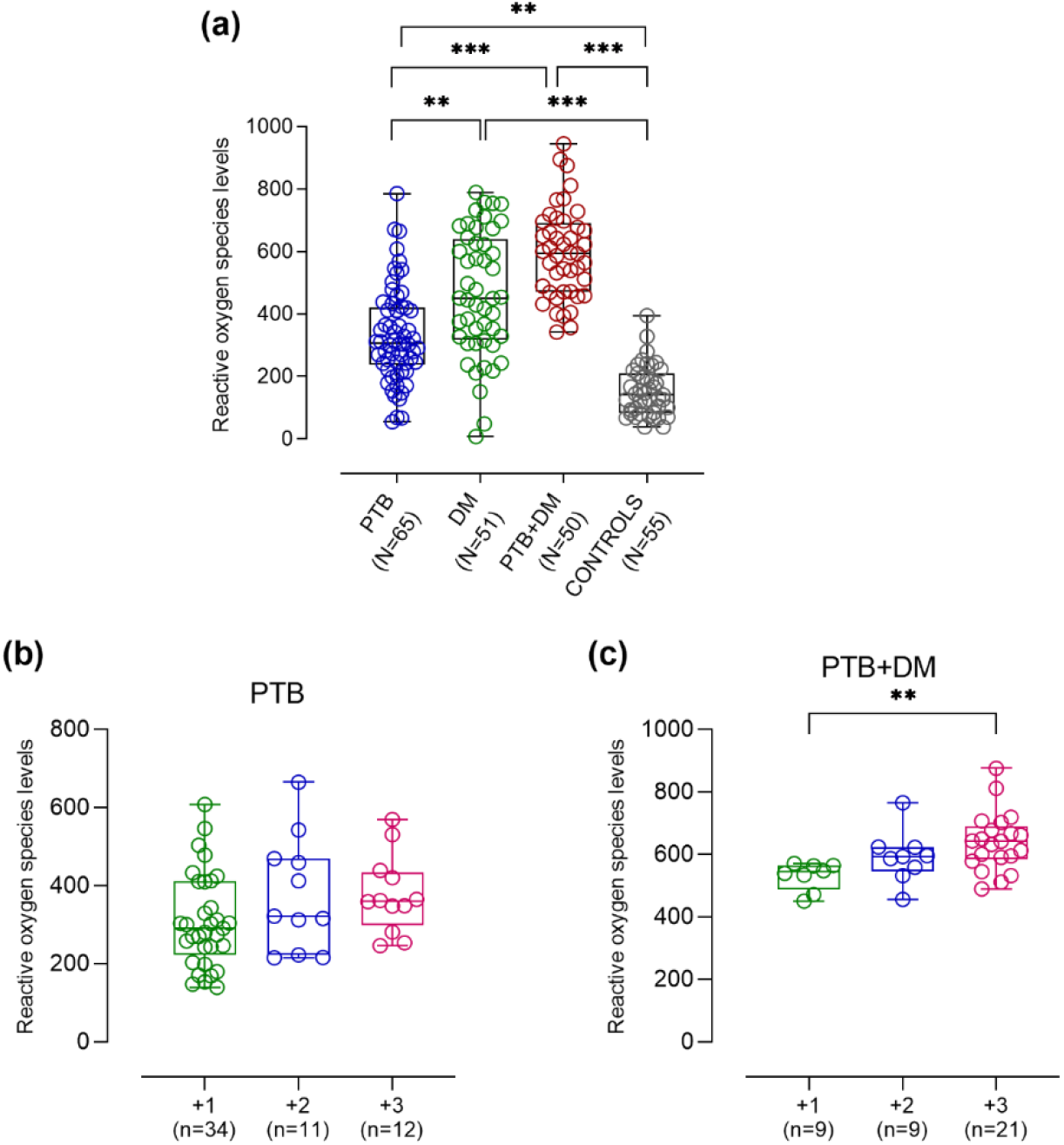
ROS levels in macrophages after BCG infection. After infection, cells were incubated with DCFDA for 30 minutes which will be converted to fluorescent compound DCF in the presence of ROS. The fluorescence signal was then read which corresponded to ROS levels. 6a shows ROS levels in PTB, DM, PTB+DM and controls. 6b and 6c shows ROS levels in different sputum grade PTB and PTB+DM patients respectively. Data is represented as fluorescent intensity and each point represents individual sample value. Box plot represents median with interquartile range. Kruskal-Wallis testing with post-hoc Dunn’s multiple comparison testing was performed. p values < 0.05 were considered to be statistically significant. One asterisk (*) indicates a p-value < 0.05; two asterisks (**) indicate a p-value < 0.01, three asterisks (***) indicate a p-value < 0.001 and four asterisks (****) indicate a p-value < 0.0001. PTB = Naïve active pulmonary TB; DM= Uncontrolled diabetic patients, PTB+DM= Uncontrolled diabetic patients with pulmonary TB, control= Healthy controls with no history of TB and DM. 1+ = 1+ sputum positive PTB/PTB+DM; 2+ = 2+ sputum positive PTB/PTB+DM; 3+ = 3+ sputum positive PTB/PTB+DM.

Taking into consideration the fact that NO is an unstable molecule with half-life of less than 5 seconds, we measured nitrites (stable serum metabolites of NO) as surrogates to measure the content of NO. The levels of NO were found to be higher in PTB group (27.32 ± 8.50 μmol/L) as compared to PTB + DM (22.76 ± 5.41 μmol/L), DM (19.44 ± 4.04 μmol/L) and healthy controls (20.60 ± 2.99 μmol/L). The difference was found to be statistically significant with higher levels in PTB group as compared to DM and PTB + DM (p<0.001 and 0.01) as shown in figure 7a. NO levels were found to be significantly decreased in 3+ sputum positive patients as compared to 2+ and 1+ sputum patients of both PTB and PTB + DM (p<0.004 and 0.005 respectively) as shown in figure 7b-7c.

**Figure 7.**
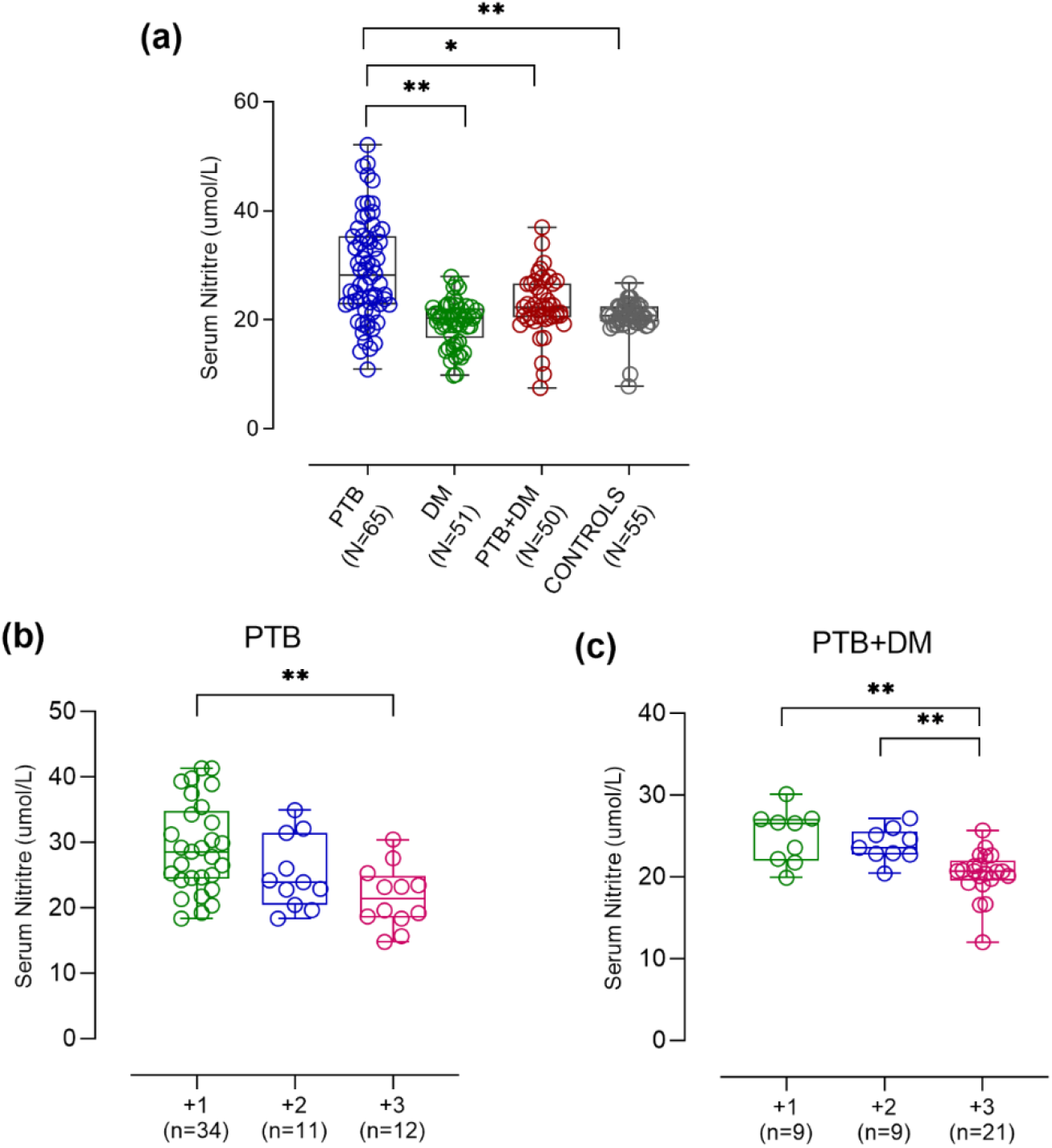
Serum NO levels in different study participants. NO was measured using Griess reaction. 7a represents NO levels in PTB, DM, PTB+DM and controls. 7b and 7c represents NO levels in different sputum grade PTB and PTB+DM patients respectively. Data is represented as median with interquartile range and each point represents individual sample value. Box plot represents median with interquartile range. Kruskal-Wallis testing with post-hoc Dunn’s multiple comparison testing was performed. p values < 0.05 were considered to be statistically significant. One asterisk (*) indicates a p-value < 0.05; two asterisks (**) indicate a p-value < 0.01, three asterisks (***) indicate a p-value < 0.001 and four asterisks (****) indicate a p-value < 0.0001. PTB = Naïve active pulmonary TB; DM= Uncontrolled diabetic patients, PTB+DM= Uncontrolled diabetic patients with pulmonary TB, control= Healthy controls with no history of TB and DM. 1+ = 1+ sputum positive PTB/PTB+DM; 2+ = 2+ sputum positive PTB/PTB+DM; 3+ = 3+ sputum positive PTB/PTB+DM.

## Discussion

Diabetics are prone to develop various infections including tuberculosis, thus increasing global TB burden. Despite its significance, the immunological and biochemical mechanisms of tuberculosis susceptibility in diabetes are not well understood. Therefore, the present study was designed to understand mechanism of increased susceptibility to TB in DM patients focusing on innate immune response. As macrophages are one of the first immune cell that mounts immune response by phagocytosing the pathogen and subsequent activation of adaptive immune response, we tried to understand any alteration in macrophage effector function under chronic hyperglycemic condition. In this case-control study, newly diagnosed pulmonary TB patients (PTB), uncontrolled type 2 diabetic patients (DM) and patients with both PTB and uncontrolled diabetes (PTB + DM) were recruited. Majority of the individuals within PTB + DM group had high bacterial burden (3+ sputum positivity) which suggests a plausible association of diabetes with increased disease severity in pulmonary tuberculosis patients.

Macrophages play a critical role in phagocytosis of mycobacteria, production of chemical mediators, antigen presentation, and subsequent activation of adaptive immune responses in host-mycobacterial interactions (13). We found significantly decreased phagocytic capacity within the macrophages of PTB + DM and DM patients plausibly due to diabetes-mediated intrinsic defect in these cells. Hyperglycemia can promote direct glycosylation of various proteins, including complements, and modify their tertiary structure, which can limit bacterial opsonization and thereby impair phagocytosis (14). A subset of TB+DM patients with HbA1c levels of 7.5-8.0 showed better phagocytosis capacity as compared to others whereas patients having HbA1c levels more than 9.5 (9 in DM and 6 in TB+DM group) showed decrease in phagocytosis levels showing adverse effect of hyperglycemia on macrophage function. Similarly, the phagocytic capacity was found to be decreased with increased disease severity, again suggesting adverse effects of hyperglycemia upon infection.

Reduction in phagocytosis is often influenced by concurrent alterations in pathogen recognition receptors that could be responsible for bacterial opsonization and internalization. Therefore, we next assessed the levels of different PRRs namely, CD11b (CR3), CD14, MARCO, TLR2 and CD206 in PTB, DM, and PTB + DM patients. Few of these receptors, such as CD14 and TLR2, induce proinflammatory cascades (15), while others, such as CD206, initiate anti-inflammatory response (16), and have been shown to be altered in diabetic environment. However, till now, limited studies have been reported on assessing the levels of these PRRs in DM patients having active TB infection.

CD11b plays critical role in mycobacterial infections in terms of cell adhesion, migration and phagocytosis (17). Decreased levels of CD11b under hyperglycemic condition in our study group may have led to decreased complement-mediated opsonization of the bacteria and hence increased susceptibility to TB infection in diabetic patients.

Another receptor, CD14 has been implicated in recognition of LAM present on mycobacterial envelope. While talking in terms of CD14 functions, a very few studies have been conducted regarding the role of CD14 in mycobacterial infections, however the findings were controversial. Our findings have shown increased surface expression of CD14 in diabetic milieu which may have adverse effect upon outcome of infection as one of the study have shown that during chronic *M.Tb.* infection, CD14 knock-out mice were shown to be protected from lethality caused by lung tuberculosis. This may be due to reduction in inflammatory response which suggest that CD14 contributes to the development of a chronic inflammatory response in the lung during tuberculosis that negatively influences the outcome of the infection (18). CD14 with the help of its co-receptors can either be beneficial to the host by induction of an adequate inflammatory and immune response, or harmful to the host due to excessive inflammation and/or dissemination of the pathogen depending on the microbe and its co-receptors (19). Therefore, we studied two co-receptors of CD14 namely MARCO and TLR2 which is required for *M.Tb.* recognition.

MARCO is a phagocytic receptor which has been implicated in host defense by recognizing trehalose 6,6′-dimycolate (TDM) which mediates potent inflammatory response via its coreceptor CD14 and TLR 2 (20). MARCO-expressing macrophages show increased propensity to phagocytose more BCG than neighbouring macrophages that do not express MARCO in the splenic marginal zone. Decreased levels of MARCO observed in PTB + DM group may suggest disruption in sensing and clearing of the *M.Tb*. pathogen. Even though, CD14 levels were higher in diabetic milieu, lower levels of MARCO receptor may have led to defective recognition and hence, increased bacterial survival in diabetic milieu. Positive correlation between CD14 and MARCO in PTB and control group is suggestive of an optimal expression of both the receptor and hence, increased bacterial uptake from CD14-MARCO receptor complex which is further supported by a lack of correlation between the two PRRs in diabetic patients with or without infection suggesting dysregulation in these receptors may have led to altered bacterial uptake and clearance.

TLR2 which is another coreceptor of CD14 was found to be higher in all the patient group which may be due to the trigger from active infection as well as inflammation (DM) which in turn can trigger various innate immune responses depending upon its coreceptor as well as the ligand to which it binds. Initial higher expression of TLR 2 along with its co receptor CD14 may worsen the outcome of infection by destructive inflammation and spread of *M.Tb*. in pulmonary tuberculosis. A positive correlation between TLR2 and CD14 in PTB + DM patients suggests increased bacterial uptake from CD14-TLR 2 coreceptor rather than CD14-MARCO coreceptor (no correlation was found in PTB + DM patients). As discussed earlier, CD14-TLR 2 overstimulation may have led to dissemination of infection and hence persistence and increased severity of infection.

Mannose receptor (CD206) is another important receptor which we have studied and found that the levels were higher in diabetic milieu (PTB + DM and DM) as compared to PTB only. CD206 plays a role in immune recognition of pathogens, following antigen internalization and presentation (21). However, this receptor is expressed in M2 macrophages that show anti-inflammatory effects during infection (22). Our data showed increased expression of CD206 in hyperglycemic condition which may suggest increased *M.Tb.* uptake via CD206 receptor which in turn may help in bacterial survival by inhibiting phagosome-lysosome fusion and shifting the macrophages to M2 phenotype which has anti-inflammatory response. This may lead to persistence, bacterial multiplication and a possible spread of infection.

Upon multivariate analysis, decreased phagocytosis capacity of macrophages in diabetic milieu was found to be associated with lower levels of CD11b, CD14 and MARCO receptors which are required for internalization of the pathogen. This suggests that hyperglycemia alters the surface expression of these PRRs and hence affects the phagocytosis capacity of the macrophages. These alterations were also associated with disease severity as CD11b, CD14, MARCO and TLR 2 were decreased and CD206 was increased in 3+ sputum positive PTB + DM patients.

Once we studied phagocytosis and various PRRs of macrophages in our study groups, we then checked for the *M.tb.* killing mechanism of macrophages via levels of reactive oxygen species (ROS) and nitric oxide (NO) in macrophages. In our study group, we observed significantly higher levels of ROS in PTB + DM patients as compared to PTB only, however PTB patients had higher levels than healthy controls. Increased levels of ROS in PTB compared to controls suggests trigger from active bacterial infection to control the infection. Although ROS plays an important role in mycobacterial killing, a recent study has shown that *M.Tb*. shows resistance against ROS during the chronic or persistent stage of infection (23). The levels were much higher in diabetic milieu (DM and PTB + DM) which may have adverse effect on infected tissue as overproduction of ROS can lead to necrosis of granuloma due to increased oxidative stress in the immune cell and hence, disruption of granuloma which may lead to dissemination of the infection. Furthermore, if the ROS levels are overwhelmed by *M.Tb*. antioxidant systems, then the pathogen will continue to survive and replicate in the host (24). Therefore, it is critical to emphasise that *M.Tb.* survival is largely dependent on the quantities of ROS produced by the host immune cells. It was also evident by our findings of increased ROS levels in 3+ sputum positive PTB + DM patients suggesting increase in disease severity upon higher ROS production under hyperglycemic condition. Therefore, higher ROS levels in PTB + DM could be counterproductive in terms of bacterial killing. Mechanistically, among all the ROS and RNS, nitric oxide (NO) is known to be one of the major contributors as an anti-TB agent and it is synthesized by inducible form of nitric oxide synthase, iNOS. As expected, nitric oxide levels were higher in PTB group suggesting active infection and induction of NO via iNOS. NO can however, have a ying-yang effect on the clearance of infection and the inflammatory response depending upon the mycobacterial strain (25). In case of PTB + DM patients, decreased levels of NO under hyperglycemic condition suggests an alteration in iNOS activity as it was shown previously that high glucose condition leads to glycation of several proteins including iNOS which may affect their activity (26). Unlike ROS, NO levels were found to be significantly lower in 3+ sputum positive patients suggesting protective role of NO in *M.tb*. infection.

## Conclusion

Our findings reveal a dynamic link between diabetes mellitus and tuberculosis infection, with a complicated interplay between aetiology and pathological progression. Modulation of anti-bacterial immune responses of macrophages in terms of defective bacterial uptake and clearance under chronic hyperglycemic condition might be one of the plausible reasons for increased susceptibility of TB in diabetics individuals. The present findings open up new research questions where a detailed mechanistic study for an in-depth understanding of cellular basis of TB susceptibility in DM is required. This will further help in better understanding of the mechanisms of hyperglycaemia impairing host defences in TB and thus lead to rational development of therapeutic strategies to alleviate the dual burden of DM and TB.

## STATEMENT AND DECLARATIONS

### Acknowledgement

We thank the DOTS center, AIIMS and Safdarjung Hospital and all the study subjects for participation in the study. We thank SERB-DST for providing research grant for the study (EEQ/2017/000165)

### Availability of data and material

All the data used for making the conclusions are in the manuscript.

### Code availability

Not applicable

### Author Contribution

Archana Singh and Sudhasini Panda conceptualized and designed the study. Sudhasini Panda drafted the manuscript. Sudhasini Panda and Diravya M Seelan carried out recruitment of patients under guidance of Archana Singh, Anant Mohan, Naval K Vikram and Neeraj Kumar Gupta. Sudhasini Panda and Diravya M Seelan carried out sample collection, standardisation and execution of experimental work along with data acquisition and interpretation of data under guidance of Archana Singh, Kalpana Luthra, Jayanth Kumar, Lakshmy R. Shah Faisal carried out execution of experimental work along with data acquisition. Archana Singh critically reviewed and contributed to the final version of manuscript. Archana Singh gave the final approval of manuscript submission and supervised the project.

### Ethics Approval

The study was conducted adopting the ethical principles stated in the latest version of Helsinki Declaration as well as the applicable guidelines for good clinical practice (GCP). Ethical approval was obtained from Institutional Ethics Committee of All India Institute of Medical Sciences, New Delhi (Ref NO: IECPG-374/28.09.2017).

### Consent to participate

Informed consent was obtained from all individuals participants included in the study.

### Consent for publication

The authors affirm that human research participants provided informed consent for publication of data.

